# Audio-visual synchrony and spatial attention enhance processing of dynamic visual stimulation independently and in parallel: a frequency-tagging study

**DOI:** 10.1101/128918

**Authors:** Amra Covic, Christian Keitel, Emanuele Porcu, Erich Schröger, Matthias M Müller

**Author notes:** Joint first authors, equal contributions (Amra Covic, Christian Keitel).

## Abstract

The neural processing of a visual stimulus can be facilitated by attending to its position or by a co-occurring auditory tone. Using frequency-tagging we investigated whether facilitation by spatial attention and audio-visual synchrony rely on similar neural processes. Participants attended to one of two flickering Gabor patches (14.17 and 17 Hz) located in opposite lower visual fields. Gabor patches further “pulsed” (i.e. showed smooth spatial frequency variations) at distinct rates (3.14 and 3.63 Hz). Frequency-modulating an auditory stimulus at the pulse-rate of one of the visual stimuli established audio-visual synchrony. Flicker and pulsed stimulation elicited stimulus-locked rhythmic electrophysiological brain responses that allowed tracking the neural processing of simultaneously presented stimuli. These steady-state responses (SSRs) were quantified in the spectral domain to examine visual stimulus processing under conditions of synchronous vs. asynchronous tone presentation and when respective stimulus positions were attended vs. unattended. Strikingly, unique patterns of effects on pulse- and flicker driven SSRs indicated that spatial attention and audiovisual synchrony facilitated early visual processing in parallel and via different cortical processes. We found attention effects to resemble the classical top-down gain effect facilitating both, flicker and pulse-driven SSRs. Audio-visual synchrony, in turn, only amplified synchrony-producing stimulus aspects (i.e. pulse-driven SSRs) possibly highlighting the role of temporally co-occurring sights and sounds in bottom-up multisensory integration.

## 1. INTRODUCTION

Behavioral goals, as well as the physical properties of sensory experiences, shape how neural processes organize the continuous and often rich influx of sensory information into meaningful units. One such process, selective attention, serves to prioritize currently behaviorally relevant sensory input while attenuating irrelevant aspects (Posner et al., 1980; Treisman and Gelade, 1980). In a visual search display, for example, items matching the color or orientation of a pre-defined target stimulus undergo prioritized processing relative to other items (Treisman and Gelade, 1980; Wolfe, 1994; Wolfe et al., 1989).

Another process exploits the spatial and temporal structure of dynamic sensory input, extracting regularities either in the visual modality alone (Alvarez and Oliva, 2009; Lee, 1999) or, by cross-referencing co-occurrences across sensory modalities (Fujisaki and Nishida, 2005). In fact, aforementioned visual search can be drastically improved by presenting a spatially uninformative tone pip that coincides (repeatedly) with a sudden change in target appearance in a dynamic search array (Van der Burg et al., 2008).

This pop-out effect has been ascribed to a gain in relative salience of the target stimulus caused by the unique integration of auditory and visual information. The impression of a multisensory object hereby hinges on the temporal precision of coinciding unisensory inputs, also termed audio-visual synchrony, a critical cue for multisensory integration (Werner and Noppeney, 2011). Consecutive synchronous co-occurrences of the same auditory and visual stimulus components further increase the likelihood of multisensory integration (Parise, 2012).

Generalizing this multisensory effect to our everyday experience of dynamic cluttered visual scenes, Talsma et al (2010) put forward that multisensory objects tend to involuntarily attract attention towards their position. As a consequence, they would gain an automatic processing advantage over unisensory stimuli. In a task that requires a sustained focus of attention on a specific position in the visual field multisensory stimuli may then act as strong distractors (Krause et al., 2012) because they withdraw common processing resources from the task-relevant focus of attention.

Interestingly, this influence seems to work both ways: As Alsius et al. (2005) have shown focusing on a visual task impedes the integration of concurrent but irrelevant visual and auditory input. This effect has been related to the concept of the temporal binding window, a period during which co-occurring attended visual and auditory stimuli are most likely to be integrated (Colonius and Diederich, 2012). The window can expand for stimuli appearing at attended locations but remains unaffected (or contracts) when spatial attention is averted (Donohue et al., 2015).

Both phenomena - the involuntary orientation of spatial attention towards multisensory events as well as impeded multisensory integration when maintaining focused attention - have largely been studied in isolation (Talsma et al., 2010). We frequently encounter situations, however, in which the two biases can act concurrently. Moreover, they may fluctuate between having conjoined and conflicting effects depending on whether attended positions and multisensory events overlap or diverge in the visual field (that is in addition to their own inherent temporal variability (Keil et al., 2012).

This complex interplay therefore warranted a dedicated investigation in a paradigm that allowed contrasting both cases directly. In the present study, we manipulated trial by trial whether participants attended to a dynamic audio-visual synchronous stimulus while leaving a concurrently presented asynchronous stimulus unattended or vice versa.

We probed early cortical visual processing by tagging stimuli with distinct temporal frequencies (Norcia et al., 2015; Regan, 1989). This frequency-tagged stimulation elicited periodic brain responses, termed steady-state responses (SSRs). SSRs index continuous processing of individual stimuli in multi-element displays and have been demonstrated to indicate the allocation of spatial attention (Kim et al., 2007; Müller et al., 1998a; Walter et al., 2012) as well as audio-visual synchrony (Jenkins et al., 2011; Keitel and Müller, 2015; Nozaradan et al., 2012).

Crucially, employing frequency-tagging allowed us to tease apart the relative facilitating effects of both factors as follows: Our paradigm featured two Gabor patches, one per lower visual hemifield, that each displayed two rhythmic physical modulations: As in classical frequency-tagging experiments they displayed a simple on-off flicker at different rates (14.17 and 17 Hz, respectively). Additionally, spatial frequencies of the Gabor patches modulated at slower rates (3.14 and 3.62 Hz, respectively), which gave the impression of a pulsation-like movement (see *Figure 1*). We exploited this pulsation to introduce audio-visual synchrony with a concurrently presented tone that carried a frequency modulation with the same temporal profile as one of the visual stimulus’ movement (Giani et al., 2012; Hertz and Amedi, 2010 for similar approaches; see Keitel and Müller, 2015). Participants were then cued randomly on each trial to attend to one of the two stimulus positions, while one of the two Gabor patches pulsed in synchrony with the tone. This paradigm enabled comparisons of SSR-indexed visual processing between four cases of Gabor patch presentation: attended synchronous (*A+S+*), attended asynchronous (*A+S-*), unattended synchronous (*A-S+*) and unattended asynchronous (*A-S-*).

**Figure 1.**
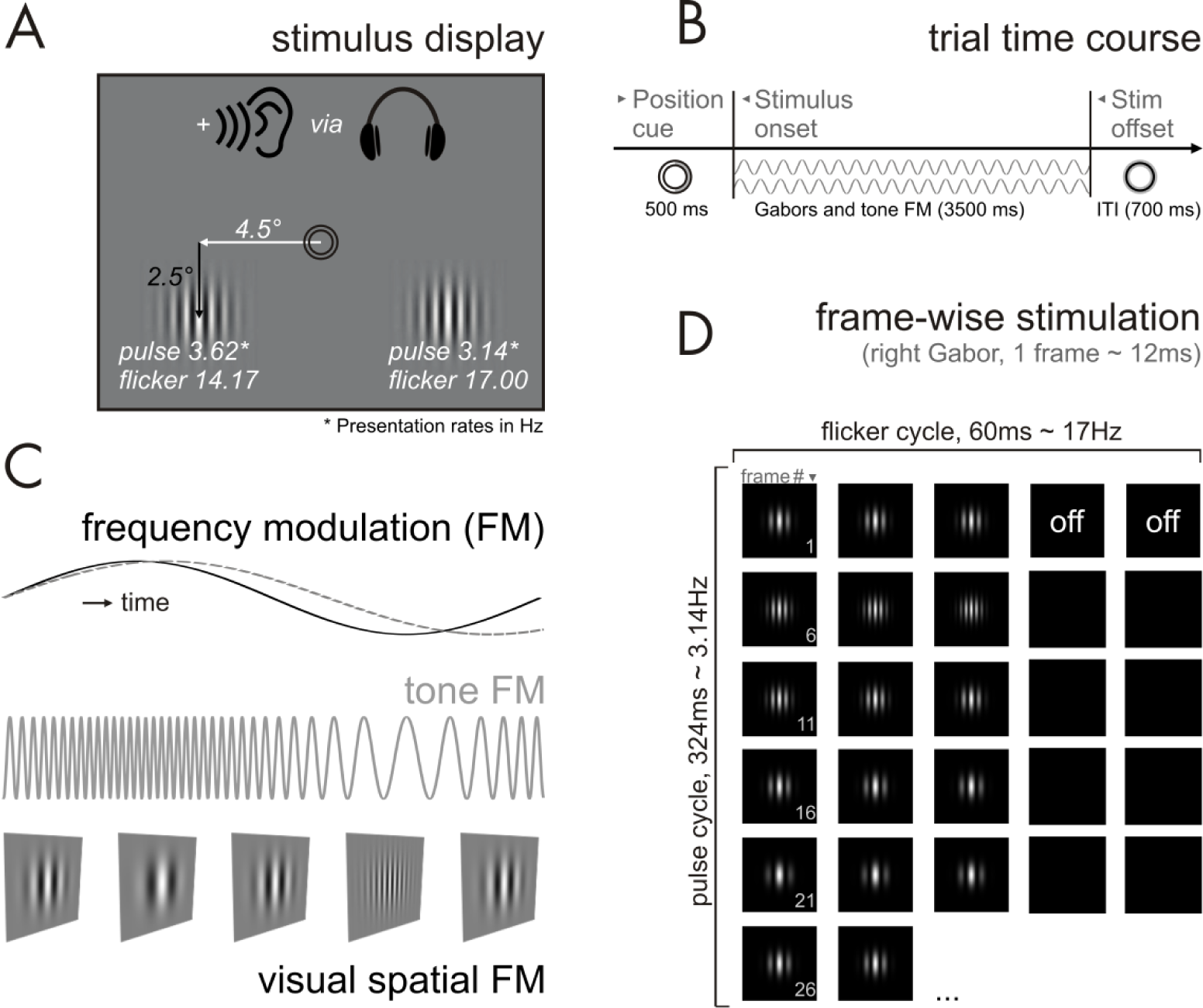
Stimulation details. (A) On-screen stimulus display comprising central fixation rings and one Gabor patch per lower left and right visual hemifield. All items not to scale. Participants received auditory stimulation via headphones. (B) Schematic trial time course. An instructive position cue allocates attention to the left or right stimulus. Subsequent ongoing Gabor-patch and tone stimulation are represented by grey sinusoids. (C) A common frequency modulation (FM; solid black line) of auditory tone pitch and the spatial frequency of one of the two Gabor patches produces a synchronous pulsing audio-visual percept. Concurrently, the spatial frequency of the other Gabor patch modulates at a slightly different frequency (dashed grey line), thus rendering it asynchronous to the tone. (D) Frame-by-frame visual stimulation for the right Gabor patch. The illustration shows the first 27 frames of each trial. Note the emphasis on the on–off cycles leading to a 17-Hz flicker along the horizontal axis (black boxes = off-frames) and one full cycle of the spatial frequency modulation leading to a 3.14-Hz ‘pulsation’ along the vertical axis.

We expected our data to replicate well-described gain effects of top-down cued spatial attention on flicker-driven SSRs (Keitel et al., 2013; Kim et al., 2007; Müller et al., 1998a). Further, we assumed that these gain effects extend to pulsation-driven SSRs, because spatial attention should prioritize any information presented at an attended location.

Secondly, we hypothesized that in line with previous findings (Nozaradan et al., 2012) audio-visual synchrony produced gain effects on SSRs. In contrast to attentional gain, results of an earlier investigation suggested that synchrony-related gain effects may be specific to pulsation-driven SSRs. Using a paradigm similar to the present study, Keitel and Müller (2015) found that an SSR component with a frequency of twice the pulsation rate was exclusively susceptible to synchrony-related gain effects. At this rate, the stimulation presumably contained strong transients critical for establishing audio-visual synchrony (Werner and Noppeney, 2011). If that were the case the current paradigm was expected to produce similarly selective effects. Alternatively, however, if audio-visual synchrony simply attracted spatial attention, then synchrony-related facilitation should mirror the pattern of attention-related gain effects on pulse- and flicker-driven SSRs. More specifically, synchrony alone should produce gain effects for flicker-driven SSRs.

Comparable patterns of attention- and synchrony-related facilitation would further point towards an account in which they may draw upon similar resources and therefore interact in facilitating visual processing: An attended stimulus would benefit less from audio-visual synchrony compared with an unattended synchronous stimulus, because attention has already been allocated to its position. Conversely, if attention- and synchrony-related facilitation relied on distinct neural resources, they were assumed to have independent additive effects on SSRs.

The latter finding could then be cast in a framework in which spatial attention biases are conveyed top-down via a fronto-parietal cortical network (Corbetta and Shulman, 2002), whereas audio-visual synchrony may have been established bottom-up via direct cortico-cortical connections or subcortical relays (Lakatos et al., 2009; van Atteveldt et al., 2014).

## 2. METHODS

### 2.1. Participants

We collected data from 14 participants with normal or corrected-to-normal vision and normal hearing. Participants gave informed written consent prior to experiments. None reported a history of neurological diseases or injury. They received course credit or a small monetary compensation for participation. The experiment was conducted in accordance with the Declaration of Helsinki and the guidelines of the ethics committee of the University of Leipzig.

Two participants showed excessive eye movements during EEG recordings and were thus excluded. Data of 12 participants aged 18 – 31 years (all right-handed, 9 female) entered analyses. Previous studies have used comparable sample sizes to reliably (re)produce effects of spatial attention (Ding, 2005; Müller et al., 1998a; 1998b; Walter et al., 2015; Zhang et al., 2010) and audio-visual synchrony (Jenkins et al., 2011; Keitel and Müller, 2015; Nozaradan et al., 2012) on SSRs.

### 2.2. Stimulation

Stimuli were presented on a 19-inch cathode ray tube screen positioned 0.8 m in front of participants. The screen was set to a refresh rate of 85 frames per second and a resolution of 1024 x 768 pixel (*width* x *height*). Visual experimental stimulation consisted of two monochrome Gabor patches with a diameter of ∼3° of visual angle, one located in the lower left and the other one located in the lower right visual field at eccentricities of 4.5° from vertical and 2.5° from horizontal meridians (see *Figure 1a*). Stimuli were presented against a grey background (RGB: 128,128,128; luminance = 30 cd/m^2^). Two black concentric circles (.4° of visual angle outer eccentricity, RGB: 0, 0, 0) in the center of the display served as fixation point.

Both Gabor stimuli underwent two independent periodic changes in the course of a trial: (1) The right patch presentation followed a cycle of 4 on-frames and 2 off-frames (2/1 on/off-ratio) resulting in a 17 Hz flicker. The left patch flickered at a rate of 14.2 Hz achieved by repetitive cycles of 3 on-frames and 2 off-frames (3/2 on/off-ratio). (2) While flickering, the spatial frequency of the Gabor patches oscillated between a maximum of 2 Hz/° and a minimum of 1 Hz/° at a rate of 3.14 Hz for the right patch and 3.62 Hz for the left patch. Periodic spatial frequency changes gave the impression of alternating contractions and relaxations that led to the percept of pulsing Gabor patches over time (*Figure 1c & d*). Pulse frequencies were chosen based on pilot experiments that served to determine a trade-off frequency range in which pulsing was readily perceptible, yet, still allowed driving periodic frequency-following brain responses (SSRs).

In addition to the visual stimuli we presented a tone with a center frequency of 440 Hz binaurally via headphones. The frequency of the tone was rhythmically modulated following sinusoidal excursions from the center frequency (10% maximum excursion = ±44 Hz). On each trial the modulation rate exactly matched the pulse rate of one of the two Gabor patches. Common rhythmic changes over time resulted in sustained audio-visual synchrony (see e.g. Schall et al., 2009).

Prior to the experiment, we employed the method of limits (Leek, 2001) to approximate individual hearing thresholds using one of the experimental stimuli, a 3.14-Hz frequency modulated tone (see e.g. Herrmann et al., 2014; Keitel and Müller, 2015). In our implementation, participants listened to a series of 10 tone sequences with a maximum duration of 15 s per sequence. Tone intensity changed during each sequence while alternating between log-linear decreases and increases across sequences. Participants were instructed to indicate by button press when they stopped or started hearing respective tones. Cross-referencing button response times with tone intensity functions yielded individual estimates of psychophysical hearing thresholds, i.e. sensation levels (SL). In the experiment, acoustical stimulation was presented at an intensity of +35 dB SL.

### 2.3. Procedure and Task

Participants were seated comfortably in an acoustically dampened and electromagnetically shielded room and directed gaze towards the fixation ring on the computer screen. At the beginning of each trial, participants were cued to attend exclusively to the left or the right visual stimulus. To this end, a green semi-circle appeared inside the fixation ring for 500 ms to indicate the task-relevant Gabor patch (see *Figure 1b*). Subsequently, the two pulsing Gabor patches and the tone were presented for 3500 ms. At the end of each trial, the fixation ring remained on screen for an extra 700 ms allowing participants to blink before the next trial started.

Participants were instructed to respond to occasionally occurring luminance changes of the cued Gabor patch (= targets) while ignoring similar events in the other patch (= distractors). During such events, Gabor patch luminance faded out to a minimum of 50% and back in within a 300 ms interval. Targets and distractors occurred in 50% of trials and up to 3 times in one trial with a minimum interval of 800 ms between subsequent onsets. Behavioral responses were recorded as space-bar presses on a standard keyboard. The responding hand was changed halfway through the experiment with the starting hand counterbalanced across participants.

We manipulated the two factors *attended position* (left vs. right Gabor patch) and audio-visual *synchrony* between attended Gabor patch and tone (synchronous vs. asynchronous) in a fully balanced design. Trials of the resulting four conditions – attended synchronous (*A+S+*), attended asynchronous (*A+S-*), unattended synchronous (*A-S+*) and unattended asynchronous (*A-S-*) – were presented in a pseudo-randomized order. Note that the tone was always in sync with one of the two Gabor patches. Therefore, in the two conditions in which the tone was out of sync with the attended Gabor patch, it was in sync with the unattended patch.

In total, we presented 600 trials (= 150 trials per condition) divided into 10 blocks (∼5 min each). Before the experiment, participants performed training for at least one block. After each training and experimental block, they received feedback on the average hit rate and reaction time.

### 2.4. Behavioral data recording and analyses

Responses were considered a ‘hit’ when the space bar was pressed between 200 to 1000 ms after target onset. We further defined false alarms as responses to distractors within the same time range. Based on these data, we calculated the response accuracy as the ratio of correct responses to the total number of targets and distractors for each condition and participant as follows:

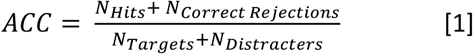

where correct responses (= numerator) are the sum of target hits *N_Hits_* and correctly rejected distracters *N_Correct Rejections_*. Correct rejections were defined as the total number of presented distracters minus the number of false alarms. Accuracies were subjected to a two-way repeated measures analysis of variances (ANOVA) with factors of *attended position* (left vs. right Gabor patch) and *synchrony* (synchronous vs. asynchronous). Response speed, quantified as median reaction times, was analyzed accordingly.

For all repeated measures ANOVAs conducted in this study effect sizes are given as *η^2^* (eta-squared). Where applicable, the Greenhouse–Geisser (GG) adjustment of degrees of freedom was applied to control for violations of sphericity (Greenhouse and Geisser, 1959). Original degrees of freedom, corrected p-values (*P*_GG_) and the correction coefficient epsilon (ε_GG_) are reported.

Further Post-hoc tests – two-tailed t-tests for paired comparisons or against zero – were applied where necessary. We applied the Holm-Bonferroni procedure to correct *p*-values (*P*_HB_) for multiple comparisons (Holm, 1979).

### 2.5. Electrophysiological data recording

EEG was recorded from 64 scalp electrodes that were mounted in an elastic cap using a BioSemi ActiveTwo system (BioSemi, Amsterdam, Netherlands) set to a sampling rate of 256 Hz. Lateral eye movements were monitored with a bipolar outer canthus montage (horizontal electrooculogram). Vertical eye movements and blinks were monitored with a bipolar montage positioned below and above the right eye (vertical electrooculogram). From continuous data, we extracted epochs of 3500 ms starting at audio-visual stimulus onset. In further preprocessing, we excluded 50% of epochs per condition (= 75) that corresponded to trials containing transient targets and distractors (= brief luminance fadings). These contained neural activity caused by processing target stimuli or motor activity due to response button presses that may have biased spectral estimates. Epochs with horizontal and vertical eye movements exceeding 25 μV (= 2.5° of visual angle), or containing blinks were also discarded. To correct for additional artefacts, such as single noisy electrodes, we applied the ‘fully automated statistical thresholding for EEG artefact rejection’ (Nolan et al., 2010). This procedure corrected or removed epochs with residual artefacts based on statistical parameters of the data. Artefact correction employed a spherical-spline-based channel interpolation. For each participant FASTER interpolated up to 4 electrodes (median = 2) across recordings and an average of up to 5.6 electrodes (minimum = 1.9, median = 3.6) per epoch. Note that epochs with more than 12 artefact-contaminated electrodes were excluded from further analysis. In total, we discarded an average of 15% of epochs per participant and condition. Subsequently, data were re-referenced to average reference and averaged across epochs for each condition and participant, separately. Basic data processing steps such as extraction of epochs from continuous recordings and re-referencing made use of EEGLAB (Delorme and Makeig, 2004) in combination with custom routines written in MATLAB (The Mathworks, Natick, MA).

### 2.6. Electrophysiological data analyses

In our analyses we focused on two neural markers that have been repeatedly demonstrated to index attentional modulation: SSR amplitudes (Morgan et al., 1996; Müller and Hubner, 2002; Quigley and Müller, 2014) and SSR inter-trial phase coherence (ITC, Kashiwase et al., 2012; Kim et al., 2007; Porcu et al., 2013). Both measures also reflect effects of audio-visual synchrony on early visual processing (Nozaradan et al., 2012). Approaches to derive amplitudes and inter-trial phase coherence differ slightly and are thus described separately below. Both approaches required spectral decompositions of EEG time series for which we used the Fieldtrip toolbox (Oostenveld et al., 2011).

#### 2.6.1. SSR power

Artefact-free epochs were truncated to segments of 3000 ms that started 500 ms after audio-visual stimulation onset and averaged separately for each EEG sensor, experimental condition and participant. The first 500 ms were omitted in order to exclude event-related potentials to stimulus onset from spectral analyses. From de-trended (i.e. linear trend removed) 3000 ms segments we quantified power (= squared amplitude) spectra by means of Fourier transforms. For the FFT, the 768 data points representing each 3000 ms segment were zero-padded to a length of 8192 (2^13) to achieve a fine-grained spectral resolution (0.0312 Hz).

*Figure 2a* illustrates that our stimulation was effective in driving distinct SSRs: Power spectra pooled across all 64 scalp electrodes and experimental conditions showed clear peaks at the stimulation rates. Notably, spectra revealed strong harmonic responses at twice the pulse frequencies (6.28 and 7.24 Hz). We included these pulse-driven harmonics in further analyses because fundamental and harmonic responses have been hypothesized to reflect different aspects of stimulus processing (Kim et al., 2011; Pastor et al., 2007; Porcu et al., 2013) and showed modulation by synchrony in a previous study (Keitel and Müller, 2015). Grand-average topographical distribution of pulse-driven as well as flicker-driven SSR power averaged over conditions showed widespread maxima at parieto-occipital electrode sites (scalp maps in *Figure 2a*) that are typically observed in experiments with lateralized flicker stimulation (see e.g. Keitel et al., 2013).

**Figure 2.**
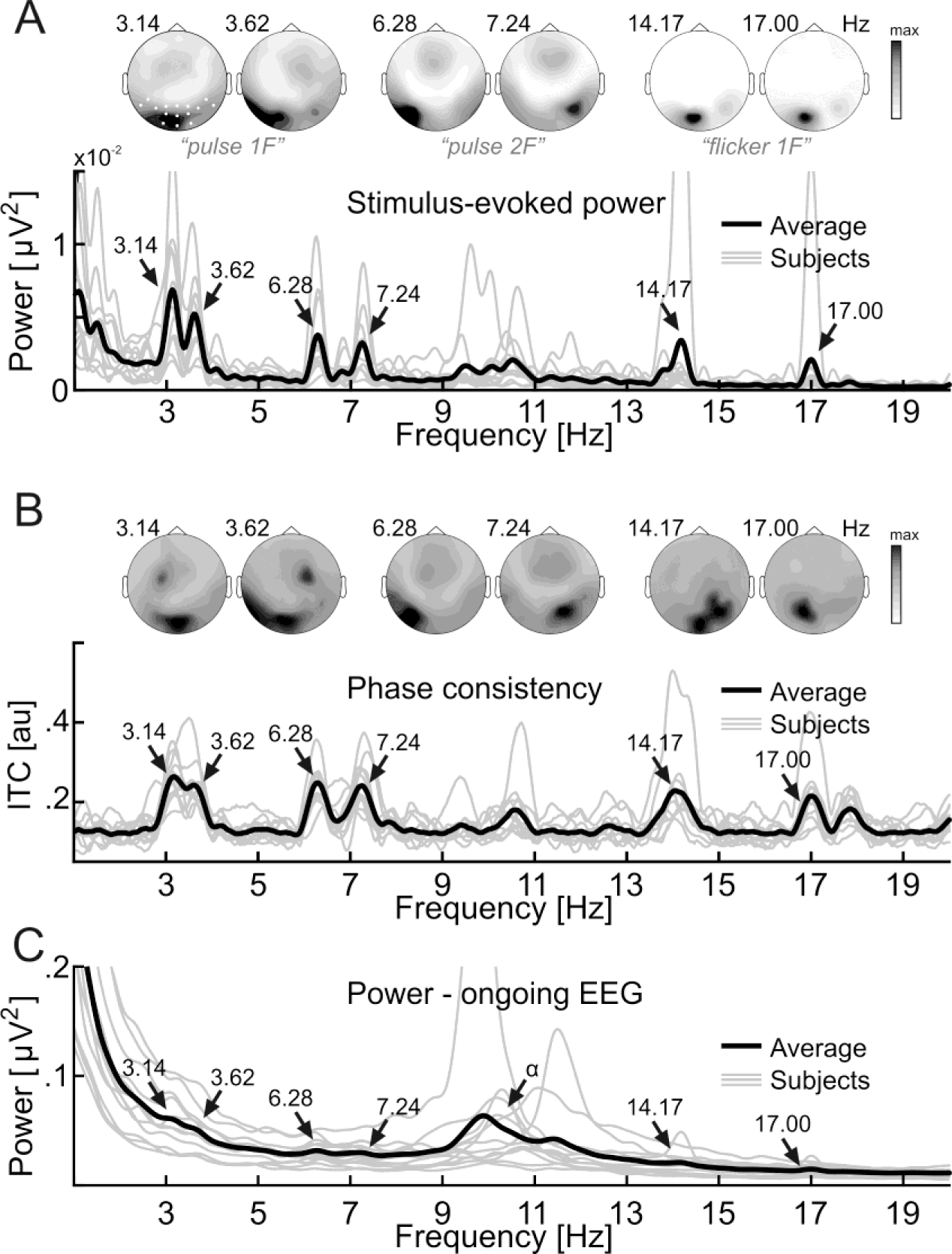
Stimulus-driven steady-state responses (SSRs) – spectra and scalp maps. (A) SSR power extracted from spectral decomposition of trial-averaged EEG waveforms, thus “stimulus-evoked”. Scalp maps show topographical distributions of power for the pulse-frequency following (*pulse 1f*), pulse-frequency doubling (*pulse 2f*) and flicker-frequency following (*flicker 1f*) SSR components driven by left and right stimuli respectively. White dots in left-most scalp map highlight the uniform sensor cluster used in all data analyses. Spectra below depict condition-averaged individual power spectra (grey lines) and, superimposed in black, the grand-average spectrum. Arrows indicate peaks that correspond to the respective driving frequencies (in Hz). (B) Same as (A) but for SSR inter-trial phase consistency (ITC) measured in arbitrary units (au). (C) Power spectra based on averaged spectral decompositions of single trials for comparison. Note that this approach emphasizes spectral characteristics of the ongoing EEG, such as the alpha rhythm (see peaks around 10 Hz, denoted *α*), over SSRs given our stimulation.

For each participant and condition, SSR amplitudes were averaged across a cluster of 15 electrodes covering parieto-occipital maxima (Oz, O1, O2, Iz, I1, I2, POz, PO3, PO4, PO7, PO8, P7, P8, P9, P10; as indicated in left-most scalp map in *Figure 2a*). Using a unified cluster of electrodes across frequencies & stimuli allowed for a comparable spatial sampling of all SSR components.

Amplitudes were further normalized by taking the decadic logarithm, then multiplying it by 20, to yield dB-scaled values (termed log-power in the following). All-positive SSR amplitude values typically show a left-skewed distribution across participants. By taking their logarithm we approximated a normal distribution (skew minimized) that better met the requirements of parametric statistical procedures.

SSR log power was subjected to four-way repeated measures analysis of variances (ANOVAs) with factors of driving *stimulus position* (left vs. right hemifield), *attention* (attended vs. unattended), *synchrony* (synchronous vs. asynchronous) and *SSR component* (pulse 1f, pulse 2f and flicker 1f).

The factor *stimulus position* had no effect on SSR log power and did not show any interaction with the other factors (see *Results*). This afforded collapsing normalized power across left and right stimuli, i.e. across pulse frequency following (‘pulse 1f’) 3.14 Hz and 3.62 Hz, pulse frequency doubling (‘pulse 2f’) 6.28 and 7.24 Hz, as well as flicker frequency following (‘flicker 1f’) 14.17 and 17.00 Hz SSRs, respectively, in subsequent analyses.

#### 2.6.2. SSR inter-trial phase coherence

We computed inter-trial phase coherence (Cohen, 2014) based on Fourier transforms of artefact-free single trial epochs, truncated to 3000 ms segments (as described above for SSR amplitude analyses) according to:

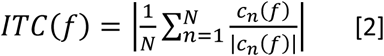

where *c_n_(f)* is the complex Fourier coefficient of trial n at frequency *f* and |.| indicates the absolute value. Inter-trial phase coherence as a measure of SSR modulation has been introduced to SSR analyses more recently (Kim et al., 2007; Nozaradan et al., 2012) and SSR amplitude and phase coherence have demonstrated different sensitivities to top-down influences on sensory processing (Kashiwase et al., 2012; Porcu et al., 2013). SSR Inter-trial phase coherence can be visualized as spectra that typically display narrow peaks at stimulation frequencies and higher order harmonics (Nozaradan et al., 2012; Ruhnau et al., 2016).

Similar to SSR amplitudes, ITCs showed broad topographic maxima at parieto-occipital electrode sites. Condition-averaged ITC spectra pooled across the 15-electrode cluster as described above (see section 2.6.1) revealed distinct peaks at the six frequencies of interest (*Figure 2b*).

Pooled ITCs were subjected to a four-way ANOVA with a design identical to SSR amplitude analyses. Note that ITCs were normalized by taking the natural logarithm prior to statistical evaluation. As for SSR log power, we found that ITC was insensitive to the *stimulus position* (left vs right; see section 3.2.2.), which again afforded collapsing across left- and right-stimulus driven in subsequent analyses.

#### 2.6.3. Power of the ongoing EEG and SSRs

As depicted in *Figure 2c*, SSRs have very low signal-to-noise ratios when being evaluated on the basis of averaged single-trial power spectra. Instead, these spectra accentuate the typical 1/f^x^ profile of power decreasing towards higher frequencies as well as peaks in the vicinity of 10 Hz that are consistent with alpha rhythmic brain activity. In turn, these features are much attenuated in SSR ‘evoked’ power and ITC spectra (*Figures 2a* and *b*).

#### 2.6.4. Joint analyses of SSR amplitude and inter-trial phase coherence modulation

As laid out in the Results section, both of our manipulations, spatial attention and audio-visual synchrony, revealed distinct patterns of effects on SSR amplitudes and ITCs. To further characterize and compare these effects we computed an index that expressed attention- and synchrony-related amplitude and ITC modulations for each subject and SSR frequency component *f* (pulse 1f, pulse 2f and flicker 1f) according to:

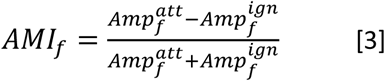

This attention modulation index (AMI) expressed the net gain effect of attention. AMIs were calculated for each stimulus individually. *Amp^att^* denotes SSR amplitudes when a stimulus was attended and *Amp^ign^* when the same stimulus was unattended (i.e. ignored). An identically scaled synchrony modulation index (SMI) was computed by contrasting SSR amplitudes between in-sync and out-of-sync conditions. We were thus able to compare both indices directly. Entering ITCs instead of SSR amplitudes into formula (3) yielded ITC-based AMIs and SMIs.

ANOVAs carried out for SSR amplitudes and ITC revealed that *attention* and *synchrony* influenced SSRs additively, i.e. no interaction between these factors was found (see *Results*). This finding justified collapsing AMIs across synchrony conditions and SMIs across attention conditions for each SSR component, separately, in the following analyses. As an example, we pooled the AMIs expressing the gain between synchronous conditions (*A+S+* vs *A-S+*) and asynchronous conditions (*A+S-* vs *A-S-*).

Because further analyses rested firmly on the assumption of an absent *attention * synchrony* interaction, we additionally applied a Bayesian inference approach because in contrast to the classical frequentist inference it allowed determining the amount of evidence in favor of the null hypothesis (*H_0_*: no interaction) explicitly. To this end, we estimated Bayes factors (Rouder et al., 2012), i.e. the plausibility of a specific model given the data. First, separately for SSR power and ITC, we determined models based on factors and interactions that turned out significant in ANOVAs. For example, SSR ITC was affected by a linear combination of factors *attention* + *synchrony* + *(synchrony * SSR component)*. These models were tested against two alternative models, one including an interaction term *(attention * synchrony)*, and another one including a main effect of *stimulus position*.

The analysis was performed by means of the function *anovaBF* provided by the R (version 3.3.0; R Core Team, 2013) package *Bayes factor* v0.9.12–2 (Morey et al., 2015). We adopted the Jeffrey-Zellner-Siow (JZS) prior with a standard scaling factor *r* of .707 (Rouder et al., 2012; 2009; Schönbrodt and Wagenmakers, 2015). Monte-Carlo resampling was based on 10^6^ iterations. Participants were considered as random factor. Importantly, Bayesian modelling favored the additive model (*attention* + *synchrony*) without an influence of the factor *stimulus position* (see *Results*) and further justified calculating AMIs and SMIs while collapsing across left and right stimuli. Results were robust against changing scaling factors. Finally, AMIs and SMIs were entered into a three-way ANOVA with factors of *SSR component* (pulse 1f, pulse 2f, and flicker 1f), *gain type* (attention vs synchrony) and *gain measure* (SSR amplitude vs ITC). Modulation indices were further tested against zero by means of t-tests (corrected for multiple comparisons).

## 3. RESULTS

### 3.1 Behavioral data

Participants detected luminance fadings more accurately when attending to left Gabor patches (main effect *attended stimulus*: F(1,11) = 32.30, *P* < 0.001, η^2^ = 0.579; see *Table 1*). Accuracy remained unaffected by in-sync vs. out-of-sync tone presentation (main effect *synchrony*: F(1,11) < 1). The interaction of both factors was not significant (F(1,11) < 1). Reaction times increased slightly when participants performed the task on in-sync Gabor patches (main effect *synchrony*: F(1,11) = 9.27, *P* < 0.05, η^2^ = 0.061; see Table 1) but were comparable between left and right stimuli (main effect *attended stimulus*: F(1,11) < 1). As for accuracy, the interaction of both factors remained negligible (F(1,11) < 1).

**Table 1.** Average behavioral performance in the visual fading detection task (N = 12).

| Attended Stimulus |  | Left |  | Right |  |
| --- | --- | --- | --- | --- | --- |
| Synchrony |  | S+ | S- | S+ | S- |
| Proportion | *M* | 85.6 % | 84.2 % | 76.4 % | 76.8 % |
| correct (%) | $\pm SEM$ | 2.2 % | 2.0 % | 2.4 % | 2.7 % |
| Reaction | *M* | 674 | 662 | 667 | 662 |
| time (ms) | $\pm SEM$ | 14 | 16 | 16 | 13 |

On average participants responded to 7.17% of distractors (median; interquartile range = 14.00%). Due to their overall low occurrence false alarms were not analysed in detail. Note however that they contributed to the here employed accuracy score (see Formula 1).

### 3.2. EEG data

We focused our analyses on SSR amplitudes and inter-trial phase coherence values (ITCs) to evaluate effects of spatial attention and audio-visual synchrony on early visual stimulus processing. Each stimulus drove three spectrally distinct SSR components: one at the frequency of stimulus pulsation, another one at twice the pulsation rate and a third following stimulus flicker (i.e., pulse 1f, pulse 2f and flicker frequencies, respectively).

#### 3.2.1. SSR power

SSR power decreased with increasing stimulus presentation rate (main effect *SSR component*: F(2,22) = 55.76, *P*_GG_ < 0.001, ε_GG_ = 0.90, η^2^ = 0.301; also see *Figure 3*) as has been documented extensively before (Keitel and Müller, 2015; Porcu et al., 2014). *Figure 3c* underlines that amplitudes further varied with the allocation of attention towards stimuli (main effect *attention*: F(1,11) = 24.15, *P* < 0.001, η^2^ = 0.094) and were affected by audio-visual *synchrony* (F(1,11) = 71.01, *P* < 0.001, η^2^ = 0.067). Amplitudes were comparable for left and right stimuli (main effect *stimulus position*: F(1,11) < 1). A significant *SSR component * synchrony* interaction (F(2,22) = 37.03, *P*_GG_ < 0.001, ε_GG_ = 0.56, η^2^ = 0.057) warranted a closer investigation of synchrony effects on specific SSR components. The crucial *attention * synchrony* interaction (F(1,11) = 1.12, *P* = 0.313, η^2^ < 0.001) as well as other interaction terms remained non-significant (maximum F(2,22) = 2.94, *P* = 0.074, η^2^ = 0.009 for the *stimulus position * SSR component interaction*).

**Figure 3.**
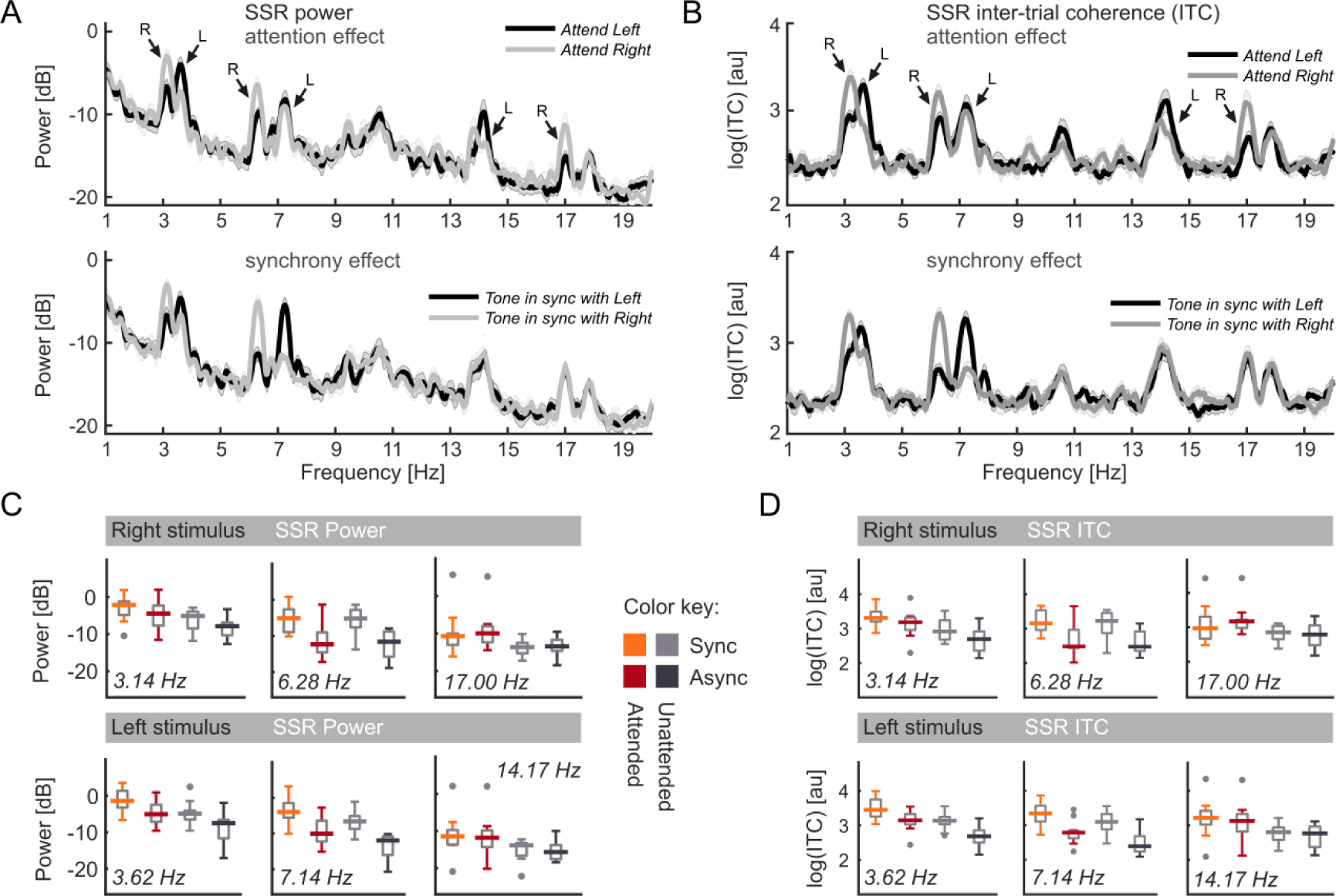
SSRs by condition. (A) Condition-resolved grand-average power (dB) spectra. Top panel: Spectra split for Attend Left (dark graph) and Attend Right (light graph) conditions. Bottom panel: Spectra split for conditions in which the tone pulsed in synchrony with the left (dark) or right (light) Gabor patch. Shaded areas represent standard error of the mean (SEM). Arrows pointing to peaks indicate the spatial position of the corresponding driving stimulus (L = left, R = right). (B) Same as in (A) but for SSR inter-trial phase coherence (ITC) measured in arbitrary units (au). (C) Zoom-in on power at SSR component frequencies. For each frequency, box plots showcase inter-individual power distributions. Boxes depict interquartile ranges with medians superimposed as strong horizontal lines. Grey dots signify outliers. A common color code applies (also see color key): Hot colors = corresponding visual stimulus attended; Monochrome = visual stimulus unattended; Light colors = visual stimulus in sync with tone; Dark colors = visual stimulus and tone asynchronous. (D) Same as in C but for SSR inter-trial coherence.

The ANOVA results suggested a model based on the linear combination of factors *attention* + *synchrony* + *SSR component* + *(synchrony * SSR component)*. Bayesian inference confirmed that this model was more plausible than the model including an *(attention * synchrony)* interaction given our data (Bf_additive_ / Bf_interactive_ = 4.61 ± 1.31%), as well as a model including a main effect of *stimulus position* (Bf_additive_ / Bf_additive + stim. pos._ = 7.55 ± 2.47%).

The *SSR component * synchrony* interaction originated from overall differences in the effect of synchrony (in-sync minus out-of-sync) on each SSR component that was most pronounced for pulse 2f components and virtually absent for flicker 1f responses (see *Figure 4a*). Specific contrasts confirmed that pulse 2f SSRs were more susceptible to synchrony effects than pulse 1f components (t(11) = 4.19, *P*_HB_ < 0.05). Pulse 1f components in turn showed stronger modulation than flicker 1f components (t(11) = 5.02, *P*_HB_ < 0.05). Lastly, pulse 2f components carried greater synchrony effects than flicker 1f components (t(11) = 7.83, *P*_HB_ < 0.05).

**Figure 4.**
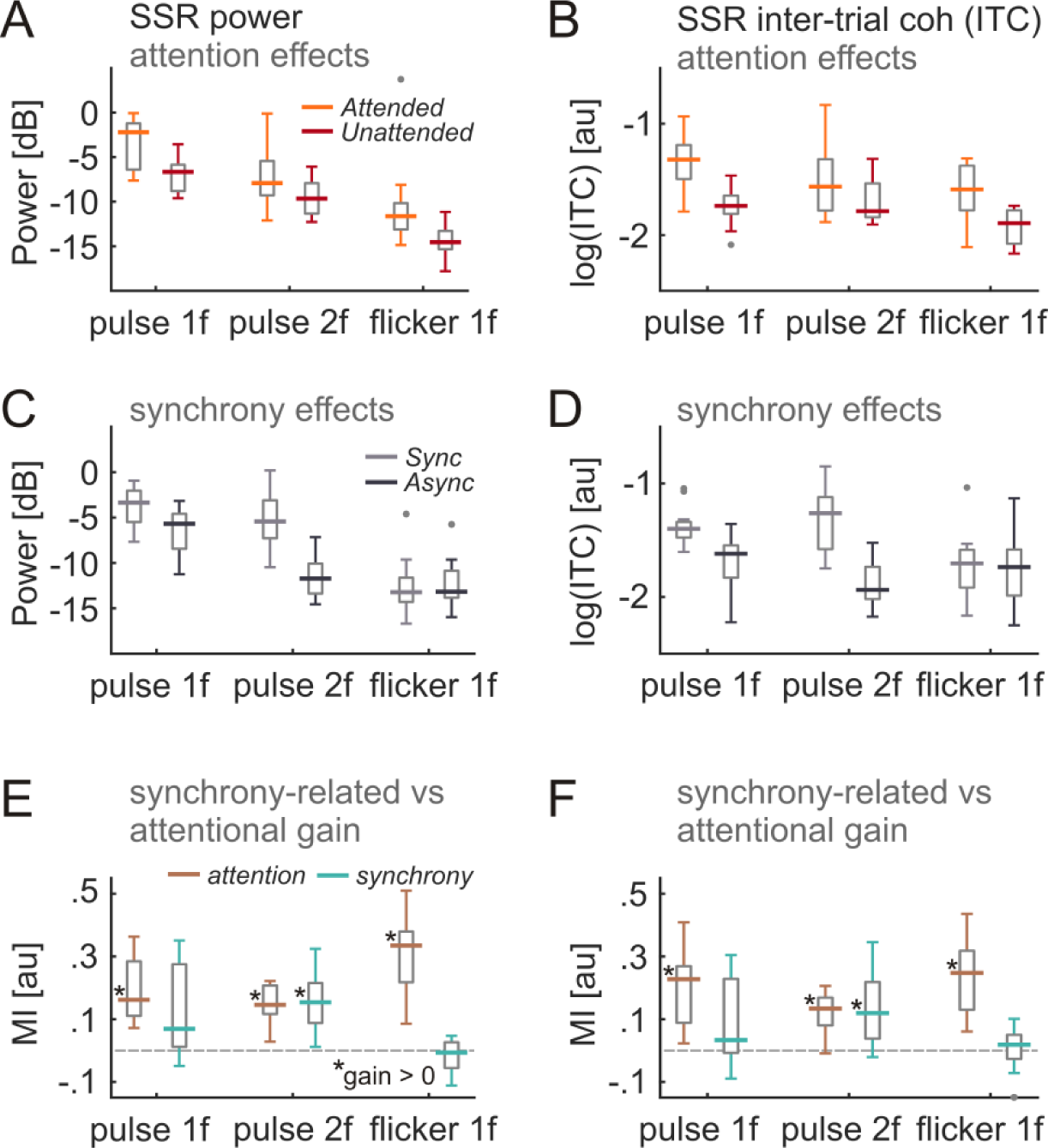
Quantifying and comparing attention- and synchrony related gain modulation. (A) SSR power (in dB) for all three SSR components of interest (*pulse 1f*, *pulse 2f* and *flicker 1f*) separated by whether the driving visual stimulus was attended (orange) or unattended (red). Box plots display inter-individual power distributions. Boxes depict respective interquartile ranges with medians superimposed as strong horizontal lines. (B) Same as in (A) but for SSR inter-trial phase coherence (ITC) measured in arbitrary units (au). (C) SSR power (in dB) for *pulse 1f*, *pulse 2f* and *flicker 1f* components separated by whether the driving visual stimulus pulsed in sync with the tone (light grey) or asynchronous (dark grey). (D) Same as in (C) but for SSR inter-trial phase coherence (ITC) measured in arbitrary units (au). (E) Boxes indicate SSR power modulation (in au) by attention (brown) and synchrony (blue) for *pulse 1f*, *pulse 2f* and *flicker 1f* components of interest. (F) Same as in (C) but for modulation of SSR inter-trial phase coherence (in au). Grey dots in plots signify outlier values. Asterisks close to medians in E & F demarcate statistically significant deviations from zero, i.e. systemic gain modulations (two-tailed t-tests, P < .05, Holm-Bonferroni corrected for multiple comparisons).

#### 3.2.2. SSR inter-trial phase coherence

ITC showed substantial variation with audio-visual *synchrony* (F(1,11) = 39.48, *P* < 0.001, η^2^ = 0.113) and the allocation of *attention* (F(1,11) = 23.43, *P* < 0.001, η^2^ = 0.139) but no effect of *SSR component* (F(2,22) = 2.24, *P* = 0.130, η^2^ = 0.026) or *stimulus position* (F(1,11) < 1). A significant *SSR component * synchrony* interaction (F(2,22) = 16.16, *P*_GG_ < 0.001, ε_GG_ = 0.54, η^2^ = 0.064) indicated that some SSR components were more susceptible to effects of audio-visual synchrony than others (*Figure 3b and d*). Remaining interaction terms, especially the *attention * synchrony* term (F(1,11) < 1), failed to indicate systematic effects (maximum F(1,11) = 2.80, *P* = 0.082, η^2^ = 0.014 for the *attention * SSR component interaction*). Only the *synchrony * stimulus position* interaction was significant (F(1,11) = 5.05, *P* = 0.046) but explained a negligible amount of variance in the data (η^2^ = 0.003) and was thus not further investigated. Note that the absence of effects of *SSR component*, *stimulus position* or an interaction of both factors on ITC supports a comparable spatial sampling (by averaging across a uniform cluster of 15 parieto-occipital electrodes; see *Methods)* of all SSR components.

Similar to SSR power, Bayesian inference supported the lack of an *attention * synchrony* interaction. Comparing additive and interactive models by means of the Bayesian approach showed evidence in favor of the additive model (Bf_additive_ / Bf_interactive_ = 4.30 ± 1.98%), again best modelled without an influence of the factor *stimulus position* (Bf_additive_ / Bf_additive + stim. pos._ = 6.71 ± 0.96%).

*Figure 4b* illustrates that the *SSR component * synchrony* interaction stemmed from greater synchrony effects (in-sync minus out-of-sync) on pulse 1f than flicker 1f components (t(11) = 4.50, p_HB_ < 0.05). Also, synchrony affected pulse 2f ITC more strongly than flicker 1f components (t(11) = 5.06, p_HB_ < 0.05). Effects between pulse 1f and 2f SSRs were comparable (t(11) = 2.09, p_HB_ = 0.19).

#### 3.2.3. Attention- vs Synchrony-related gain effects

As described in detail in the methods section, we computed indices that expressed SSR attention- and synchrony-related modulation of each SSR component. These modulation indices (AMIs and SMIs) allowed for a direct statistical comparison of the magnitude of attention and synchrony-related gain effects on SSR amplitudes and ITCs. As MI analyses assumed effects of attention and synchrony to be additive, further to the non-significant *attention * synchrony* interactions reported above, we estimated the plausibility of additive vs interactive models given our data by using a Bayesian approach. The estimated Bayes factors for SSR power and ITC (see sections 3.2.1. and 3.2.2.) indicated that both results were more than 4 times more likely under the additive than the interactive model. Comparing modulation indices based on SSR amplitudes (Figure 4E) and SSR inter-trial coherence (Figure 4F) revealed that, overall, attention led to stronger gain effects on SSRs than synchrony (15.7% ± 1.8 vs 13.7% ± 1.8, mean ± standard error; main effect *gain type*: F(1,11) = 28.79, *P* < 0.001, η^2^ = 0.20). Most importantly, however, this difference in gain effects varied between SSR components (interaction *gain type * SSR component*: F(2,22) = 6.66, *P_GG_* = 0.007, ε_GG_ = 0.898, η^2^ = 0.13) in the absence of a modulation of gain effects across *SSR components* alone (main effect: F(2,22) = 0.41, *P* = 0.668).

From a methodological perspective it should be noted that power-based modulation indicated a small but significantly higher gain than ITC based modulation (main effect *gain measure*: F(1,11) = 19.77, *P* < 0.001, η^2^ < 0.01), an effect that further depended on whether attention or synchrony caused the modulation (interaction *gain measure * gain type*: F(1,11) = 7.85, P = 0.017, η^2^ < 0.01).

However, we disregarded these small effects to investigate the *gain type * SSR component* interaction more closely. First, SSR amplitude and ITC-based modulation indices were tested against zero. Attention systematically modulated all SSR components (see Figures 4E & F; asterisks denote significant deviations from zero at a Holm-Bonferroni corrected alpha level of *P* < .05). Synchrony, in turn, only modulated pulse 2f, but not pulse 1f and flicker 1f responses for both, SSR power- and ITC- based modulation indices.

Given these highly similar patterns we pooled across measures. Then we tested gain differences (Attention minus Synchrony) between SSR components. Elucidating the *gain type * SSR component* interaction, gains differed more for flicker 1f than for pulse 1f SSRs (t(11) = 3.03, *P_HB_* < .05) and for pulse 2f SSRs (t(11) = 3.06, *P_HB_* < .05). In turn, gain differences were statistically comparable between pulse 1f and pulse 2f SSRs (t(11) = -0.92, *P* = .376) highlighting the exclusive role of the flicker-driven signal component.

## 4. DISCUSSION

The role of top-down attention in multisensory binding and, conversely, bottom-up multisensory influences on attentional orienting have been studied largely independent of each other (Talsma et al., 2010). The present study was designed to bridge this gap. Specifically, we studied situations in which participants attended to the position of one of two pulsing and flickering stimuli providing it with a top-down processing advantage over the other stimulus. Additionally, a tone pulsing in synchrony with either the attended or unattended stimulus was introduced to produce a strong multisensory bottom-up bias in visual processing. EEG-recorded SSRs driven by stimulus flicker and pulsation allowed us to test whether and how spatial attention and audio-visual synchrony acted, and possibly interacted, to facilitate cortical visual stimulus processing.

We evaluated two commonly used SSR measures, evoked power and inter-trial phase coherence (ITC) to quantify modulations in stimulus processing. Both measures widely agree on patterns of effects and will thus be considered jointly in the following.

Briefly summarizing the results, spatial attention facilitated pulse- *and* flicker-driven SSRs. In contrast, synchrony specifically facilitated pulse-driven SSRs only with greater effects on pulse 2f components while leaving flicker 1f components unaffected. Most importantly, attention and synchrony produced independent additive gain effects. We confirmed that, given our data, an additive model of both influences was more plausible than assuming interactive effects. These findings replicate results from an earlier study using a related paradigm. In that study we tested concurrent influences of feature-based attention and audio-visual synchrony on two spatially super-imposed Gabor patches (Keitel and Müller, 2015).

### 4.1. Spatial attention facilitates processing of all stimulus aspects

The described effects of spatial attention are in line with numerous studies demonstrating sensory gain effects on SSR-indexed cortical visual processing (Müller et al., 1998a; Störmer et al., 2014; Walter et al., 2015). Interestingly, our results show that spatial attention has comparable effects on SSRs driven by two different but simultaneous rhythmic changes in stimulus appearance: a relatively fast on-off flicker (> 14 Hz) and a slow-paced sinusoidal spatial frequency modulation (3 – 4 Hz). These results support the notion that spatial attention prioritizes all aspects of sensory information within its focus (Andersen et al., 2008; Keitel and Müller, 2015) as is central to psychological (Treisman and Gelade, 1980; Wolfe, 1994) and neurophysiological models of attention (Bundesen et al., 2015; Reynolds and Heeger, 2009).

Note that participants performed better in the visual detection task when they attended to the left stimulus. This effect could be due to a left-hemifield advantage as has been described previously for rapid serial visual presentation paradigms (Śmigasiewicz et al., 2014; Verleger et al., 2011). In turn, SSR analyses did not show differences in stimulus processing between left and right stimulus positions. It is therefore possible that the imbalance in task performance did not stem from differences in early visual processing of left and right stimuli but was introduced at a later processing stage.

### 4.2. Synchrony selectively facilitates stimulus aspects relevant for multisensory integration

Facilitation of visual processing by audio-visual synchrony has largely been studied using transient stimuli (Busse et al., 2005; Talsma et al., 2009). So far, only a few studies have demonstrated synchrony-driven effects while employing dynamic ongoing stimulation (Keitel and Müller, 2015; Nozaradan et al., 2012; Schall et al., 2009). Prolonged exposure to synchronous sensory input, however, can be a vital factor in multisensory integration because it improves the estimate of temporal correlations between visual and auditory stimuli over time (Parise and Ernst, 2016). This is important in situations with multiple concurrent stimuli (as studied here) because even unrelated visual and auditory events can occur simultaneously occasionally.

Our study corroborates this role of ongoing audio-visual synchrony. Interestingly, synchrony-related gain effects were thereby restricted to SSR components that reflected stimulus pulsing, i.e. those rhythmic modulations that produced the impression of synchrony.

Visual stimulus dynamics either matched with or differed from the spectral profile of the auditory stimulus, thus providing either maximal or minimal temporal correlation. Less intuitively, the SSR component at twice the pulsation rate (pulse 2f) showed greater synchrony modulations than the pulse-frequency following response (pulse 1f). In line with Keitel et al. (2015), who employed a stimulus with similar dynamic properties, the pulse 2f modulation was accounted for by the transients elicited by the stimulus at twice the stimulus pulsation rates during maximum up- and down-slopes of the sinusoidal modulation, or alternatively its extrema, i.e. peaks and troughs.

We propose that successive cross-modal phase resets may be the neural process underlying synchrony-related modulation of both pulse-driven components. Cross-modal phase resetting has been considered as the primary channel for multisensory interactions between early sensory cortices (Lakatos et al., 2009; van Atteveldt et al., 2014). Unlike neurons in higher order cortices, which are intrinsically multisensory (and hence sensitive to combined multisensory information) neurons in early sensory cortices are primarily sensory specific, but crucially sensitive to temporal information conveyed also by non-specific modalities. As underlined by Lakatos et al. (2008), appropriately timed inputs in one modality can aid in processing a stimulus presented in a different modality. In our case these connections may support phase stability of visual SSRs by providing a cross-modal temporal scaffold (Kayser et al., 2010; Lakatos et al., 2009). As a consequence, the temporal precision of cortical stimulus representations increases, which awards them a processing advantage (Chennu et al., 2009).

Although our results are broadly in line with Nozaradan et al. (2012), who firstly measured synchrony effects on SSRs, it is worth noting a discrepancy: In contrast to our findings the authors reported an effect on a flicker-driven SSR with a frequency of 10 Hz, while establishing synchrony with auditory beats at either 2.1 or 2.4 Hz. These differences may be accounted for by the fact that the authors presented only one visual stimulus centrally. In this setup, gain effects cannot not unambiguously be ascribed to synchrony, or alternatively, altered attentional demands between synchronous and asynchronous conditions.

### 4.3. Facilitatory effects of spatial attention and synchrony add up

We found that attended and unattended stimulus experienced comparable gain through synchrony. Vice versa, synchronous and asynchronous stimuli were similarly facilitated when their position was attended. Remarkably, these findings point towards a dual reign of attention and audio-visual synchrony in early sensory cortices, suggesting that both influences can work independently and in parallel. This result seemingly contradicts previous studies (Alsius et al., 2005; Fairhall and Macaluso, 2009) that showed an interdependence between attention and multisensory interactions. However, this contradiction can be reconciled by examining the experimental paradigm employed in the current study.

Unlike previous experiments, in which mutual input from different senses was essential for successful behavioral performance, it is hard to construe a direct benefit from audio-visual synchrony in performing our task, i.e. the purely visual detection of luminance changes. Our paradigm might thus have promoted the independence between attention and audio-visual interactions triggering two concurrent, but distinct processes: On the one hand, performing the detection task required a sustained goal-driven deployment of spatial attention, while on the other hand merging the audio-visual signals was most likely a stimulus-driven process, triggered by the high temporal correlation between auditory and visual signal components.

For these two processes to co-occur independently, we assumed the involvement of distinct neural pathways. Various aspects of attention and its influence on perception have been related to a number of anatomical networks (Shipp, 2004). To date, a dorsal fronto-parietal network, which entails the intra-parietal sulcus (IPS) in posterior parietal cortex, a portion of the precentral supplemental motor area, the so-called frontal eye fields (FEF) and early sensory areas, such as visual cortex has been described most comprehensively (Corbetta and Shulman, 2002). This cortical network has been implicated in the control of attention (Corbetta et al., 1998) and was likely involved in deploying the resources necessary to perform in our behavioral task.

On the other side, auditory influences on visual processing could have been conveyed by two candidate routes that have been suggested as a results of earlier invasive electrophysiological and anatomical studies in the animal brain: (1) feed-forward projections between thalamus and early sensory cortices (Cappe et al., 2009), (2) lateral projections between early sensory cortices (Falchier et al., 2002). From our data alone, we cannot say which pathway was critical in the investigated situation. Both neural pathways however are anatomically distinct from the fronto-parietal attention network (as described above) and are thus consistent with our results.

It should be mentioned that our data analyses and interpretation of results depend on the implicit assumption that attention and synchrony effects follow similar time courses and, once established, remain constant through the course of each trial. At least for, spatial attention we know that gain effects reach asymptote after ∼500 ms and keep level for several seconds (Müller et al., 1998b). A time course for synchrony-related gain instead has not been established yet. This uncertainty notwithstanding, we restricted our analyses to a period starting 500 ms after stimulus onset. We were confident that this time frame would allow for enough audio-visual coincidence to be detected to establish synchrony. The comparison of temporal profiles of attention- and synchrony related gain remains an interesting subject for future studies, nevertheless.

As a final remark, Talsma et al. (2010) suggested that bottom-up multisensory integration benefits a given stimulus the most when competition within one sensory modality is high, e.g. when the visual field is cluttered. Our situation, with one stimulus presented to each hemifield, promoted only minimal competition. Inter-hemispheric competition is introduced relatively late in the visual processing hierarchy (Schwartz et al., 2007). Moreover, attentional resources seem to split more readily between than within visual hemifields (Franconeri et al., 2012; Störmer et al., 2013; Walter et al., 2015). It would thus be interesting to test how synchrony-related gain effects vary with the amount of competition by placing more than one stimulus within visual hemifields.

### 4.4. Conclusion

We investigated the concurrent effects of spatial attention and audio-visual synchrony on early cortical visual stimulus processing. Our paradigm allowed us to test both influences in isolation as well as their combined effects. We found that attention-related and synchrony-related facilitation add up when an audio-visual synchronous stimulus is attended. Further, attention facilitated pulse- and flicker-driven neural responses while synchrony only targeted pulse-driven responses, i.e. those coding for stimulus dynamics that were relevant for multisensory integration. Consequentially, the present results favor an account in which goal-directed sustained spatial attention and stimulus-driven audio-visual synchrony convey their influences independently via different neural processes and possibly along different neural pathways. At least for situations similar to the one studied here, this finding implies that facilitation through synchrony cannot simply be modelled as a sustained attraction of spatial attention.

## Acknowledgments

Work was supported by the Deutsche Forschungsgemeinschaft (grant no. MU972/21-1). Data presented here were recorded at the Institut für Psychologie, Universität Leipzig. The authors appreciate the assistance of Renate Zahn in data collection. Experimental stimulation was realized using Cogent Graphics developed by John Romaya at the Laboratory of Neurobiology at the Wellcome Department of Imaging Neuroscience, University College London.

## Conflict of interest

The authors declare that they have no conflict of interest.

